# The comparative and genetic methods for East European Unionidae taxonomy

**DOI:** 10.1101/390872

**Authors:** Aleksandra S. Anisimova, Alibek Abdrakhmanov, Tatyana V. Neretina, Alexey S. Kondrashov, Victor V. Bogatov

**Affiliations:** Faculty of Bioengineering and Bioinformatics, Lomonosov Moscow State University, Moscow, Russia; Department of Ecology and Evolutionary Biology, University of Michigan, Ann Arbor, MI, USA; Federal Scientific Center of the East Asia Terrestrial Biodiversity, Far Eastern Branch, Russian Academy of Sciences, Vladivostok, Russia

## Abstract

Taxonomy of freshwater mussels from family Unionidae has been ambiguous for a long time. A number of methods used for their identification, including the so-called comparative method, are based on shell morphology. However, this morphology turned out to have a high level of within-species variation, and the shape of the shell of a specimen depends strongly on its environment and conditions of growth. For these reasons, the number of species recognized by the comparative method kept increasing. We applied both the comparative morphological method and methods of molecular genetics to address the taxonomy of Unionidae. We performed the comprehensive study of 70 specimens of Unionidae mussels collected from the River Ivitza, Volga basin. The specimens represented 14 comparative species, belonging to 4 comparative genera of Unionidae: *Colletopterum, Pseudanodonta, Unio* and *Crassiana*. Sequencing of the nuclear (ITS1) and mitochondrial (COI, 16S rDNA) genetic regions revealed 5 groups with high within-group genetic homogeneity separated from each other by long genetic distances. According to the comparison with the available sequences, these groups correspond to 3 Eastern European genera and 5 species: *Anodonta anatina, Pseudanodonta complanata, Unio pictorum, Unio tumidus* and *Unio crassus*. The results obtained indicate that the comparative method is inappropriate for the taxonomic analysis of East European Unionidae.

## Introduction

By the middle of the XX century, eight species of large bivalve mussels Unionidae were recognized in North-Eastern Europe (central and northern parts of the European Russia). They belong to two genera: *Unio* Philipsson in Retzius 1788 (three species) and *Anodonta* Lamarck 1799 (five species) [1,2]. Identification of these species was based mostly on shell morphology: shell shape and size of adult mussels, location of the umbo relative to the front edge of the shell, features of the umbo sculpture, thickness and convexity of the valves, tooth shape, etc.

At the same time, from the end of 1960’s the so-called comparative method has been proposed for identification of freshwater Unionidae in Soviet Union. This method is based on comparing the outlines of the cross-sections of valves by overlaying drawings on the top of each other. According to the ideas of Tompson the growth of some groups of animals, including Unionidae, follows the logarithmic spiral trajectories [3]. Based on that, the shape of the outer shell outline, perpendicular to its longitudinal axis from umbo the lower edge, was postulated to be species-specific [4–6].

By the beginning of the XXI century, as the result of usage of comparative method the number of North-Eastern European Unionidae increased to twenty seven species attributed to six genera (Table 1): *Unio* (four species); *Tumidiana* Servain 1882 (three species); *Crassiana* Servain 1882 (six species); *Pseudanodonta* Bourguignat 1876 (four species); *Colletopterum* Bourguignat 1880 (seven species); *Anodonta* (three species) [7]. By this time, some malacologists started to consider the comparative method improper for taxonomic studies of large bivalves [8,9]. According to Graf, the traditional species are regarded as a better description of actual diversity, but further revision is required [10]. The number of East European Unionidae species has been reduced, to only six species belonging to three genera (Table 1), with all comparative species names declared invalid.

**Table 1.** Species composition of Eastern European Unionidae according to different systems

| Zhadin, 1952 [2] | Starobogatov, 1977 [28] | Starobogatov et al., 2004 [7] | Graf, 2007 [10] | Bogatov, Kiyashko, 2016 [15] |
| --- | --- | --- | --- | --- |
| **Subfamily Anodontinae** |  |  |  |  |
| *Anodonta anatina* (L. 1758)<br>*A. piscinalis* (Nilsson 1823)<br>*A. p. ponderosa* (Pfeiffer 1825)<br>*A. p. volgensis* (Shadin 1938) | *Pseudanodonta anatina*<br>*A. piscinalis*<br>*A. ponderosa* [= *A. p. volgensis*]<br>*A. minima* (Millet 1883) | *Colletopterum anatinum* [= *A. minima*]<br>*Co. piscinale*<br>*Co. ponderosum*<br>*Co. nilssoni* (Kuester 1842)<br>*Co. rostratum* (Rossm. 1836) | *A. anatina* [= *Co. piscinale*, *Co. nilssoni*, *Co. ponderosum*, *Co. rostratum*] | *Co. (Piscinaliana) anatinum*<br>*Co. (P.) piscinale*<br>*Co. (P.) ponderosum*<br>*Co. (P.) nilssoni*<br>*Co. (P.) rostratum* |
| *A. cygnea* (L. 1758)<br>*A. cellensis* (Schroeter 1779) | *A. cygnea* [= *A. cellensis*]<br>*A. zellensis* (Gmelin 1791)<br>*A. stagnalis* (Gmelin 1791) | *A. cygnea*<br>*A. zellensis*<br>*A. stagnalis* | *A. cygnea* [= *A. zellensis*, *A. stagnalis*] | *A. cygnea*<br>*A. zellensis*<br>*A. stagnalis* |
| **Subfamily Pseudanodontinae** |  |  |  |  |
| *A. complanata* (Rossm. 1835) | *P. complanata*<br>*P. elongata* (Holland. 1836)<br>*P. kletti* (Rossm. 1835) | *P. complanata*<br>*P. elongata*<br>*P. kletti*<br>*P. nordenskioldi* | *P. complanata* | *P. complanata* [= *P. kletti*]<br>*P. elongata* |
| **Subfamily Unioninae** |  |  |  |  |
| *Unio pictorum*<br>(L. 1758) | *U. pictorum*<br>*U. limosus*<br>(Nilsson 1822) | *U. pictorum*<br>*U. limosus*<br>*U. protractus*<br>(Lindholm 1922)<br>*U. rostratus*<br>(Lamarck 1799) [= *U. longirostris* (Rossm. 1836)] | *U. pictorum*<br>[= *U. limosus*,<br>*U. protractus*, *U. rostratus*] | *U. (Unio) pictorum*<br>[= *U. rostratus* non *U. longirostris*]<br>*U. (U.) limosus*<br>*U. (U.) protractus*<br>[= *U. muelleri* (Rossm. 1836 partim)] |
| *U. tumidus*<br>(Philippon in Retzius 1788) | *U. tumidus*<br>*U. ovalis*<br>(Montagu 1803)<br>*U. muelleri*<br>*U. longirostris*<br>(Rossm. 1836) | *Tumidiana tumida*<br>*T. conus* (Spengler 1793)<br>*T. muelleri* | *U. tumidus*<br>[= *T. tumida*,<br>*T. conus*,<br>*T. muelleri*] | *U. (Tumidiana) tumidus*<br>[= *U. ovalis* partim,<br>*U. muelleri* partim]<br>*U. (T.) longirostris*<br>*U. (T.) conus* [= *U. ovalis* partim] |
| Subfamily **Psilunioninae** |  |  |  |  |
| *U. crassus*<br>(Philippon in Retzius 1788) | *Crassiana crassa*<br>*Cr. musiva*<br>(Spengler 1793)<br>*Cr. fuscula*<br>(Rossm. 1836)<br>*Cr. nana*<br>(Lamarck 1819)<br>*Cr. irenjensis*<br>(Kolbelt 1912) | *Cr. crassa*<br>*Cr. musiva*<br>*Cr. fuscula*<br>*Cr. irgizlaica*<br>(Lindholm 1904)<br>*Cr. nana*<br>*Cr. cyprinorum*<br>(Bourg. 1882) | *U. crassus*<br>*crassus*<br>[= *Cr. musiva*, *Cr. irgizlaica*. *Cr. nana*,<br>*Cr. cyprinorum*] | *Cr. crassa*<br>*Cr. musiva*<br>[= *Cr. fuscula*, *Cr. cyprinorum*, *Cr. irgizlaica*]<br>*Cr. nana* |

Synonymous names are indicated in square brackets; subfamily names are indicated according to Bogatov and Kiyashko [15].

In the early versions of the comparative method the authors proposed to compare the contours of shells placed under the microscope in only one fixed position without taking into account the size of the mollusk [11,12]. This approach ignored the fact that the lateral surface of the posterior part of the shell of many Unionidae mussels is further moved from the commissural plane (the plane of the closure of the valves) in the process of growth, in comparison with its anterior part. Consequently, if the shell is placed perpendicular to its frontal section, the outer contour visible under the microscope shifts to the posterior edge of the shell as shell size increases. As the result, the estimation of the curvature of profiles leads to the high variation between different sized shells of the same species. The errors of comparative method led to the unjustified description of new species, which ultimately discredited the method itself [12].

The modification of comparative method (MCM) is based on the pervasive pattern of freshwater Unionidae shell morphogenesis. It consists of the comparison of the Arc of Maximal Convexity of the Valve’s Outline (AMCVO) [11–15]. This section extends from the umbo through the points maximally shifted from the commissural plane at different times of shell formation. The AMCVOs do not change with increase of the mollusks size. The revision of the East European Unionidae based on the MCM decreased the number of species to nineteen belonging to four subfamilies and five genera (Table 1) [15]. Since the species-specific forms of the contours of maximum convex of shell have not yet been confirmed genetically, the aim of this work is to estimate the taxonomic significance of the comparative species by genetic methods.

Surprisingly, no systematic analysis of molecular data that could shed light on the validity of comparative species has yet been performed. Most of the recent attempts to estimate genetic differences between the comparative species of European *Margaritifera* [9,16,17], European *Unio* [18], Far East Russian *Cristaria* [19] and *Nodularia* [20] did not involve identification of comparative species by MCM. The identification was instead based only on measurements of the height and the width of the shell or the ratio of shell convexity to its maximum height, which does not allow to determine their taxonomy reliably [12,15].

In a recent study, Klishko et. al made an attempt to identify Anodontinae species with MCM [21]. However, even in this work some inconsistencies with the original AMCVO measurements method can be found [13–15]. First, according to Figure 2 from [21] the two disparate sets of arcs were used in the analysis. This may be the result of usage of two different microscopes with different focal lengths to the valves. Second, the arcs were compared with the frontal contour instead of the contour of maximal convexity of the valves, and the points of inflection between the surface of the shell and the ligament were selected for the initial points of spiral development instead of umbo apex. Moreover, standard measurements of shells were obtained from the shells of wide growth range instead of middle growth samples suggested in the original method [15]. As the result, AMCVO measurements were claimed to be inappropriate for taxonomy of Anodontinae species. Definitely, findings of Klishko et al. provide a rationale for testing whether the MCM can be applied to a broader range of Unionidae genera.

Moreover, studies aimed at testing the comparative method by molecular genetic analyses were mostly based on only one mitochondrial cytochrome oxidase gene region (COI) [18–20]. This approach may lead to erroneous phylogenetic reconstructions due to doubly uniparental inheritance of mitichondria intrinsic to Unionidae [22–24]. For example, in some cases nuclear and mitochondrial genomes suggest different phylogenetic relationships between species in taxonomic analysis of Unionidae [25]. To overcome these difficulties the combination of both nuclear and mitochondrial data can be applied [26,27].

## Materials and methods

### Samples collection

Specimens of Unionidae mussels were collected by V. Bogatov on May 21-22 of 2016 in the lowland River Ivitza (Volga basin, Rameshkovsky District, Tver Oblast, Russia). It flows into the Medveditza River near the Ivitza village and is not subjected to anthropogenic pollution (Fig 1). The length of the Ivitza is 51 km, and the area of its basin is 397 km^2^. The width of the river in the places of the collection is 5-12 m with the depth up to 1 m and bottom soil type ranging from silt to sand. The velocity ranges from 0.05 to 0.1 m/s at the stream pools and speeds up to 0.3 m/s at the rapids. Low waters and spits can be found in the river. The Ivitza River is covered by ice from December to April [29].

**Fig 1.**
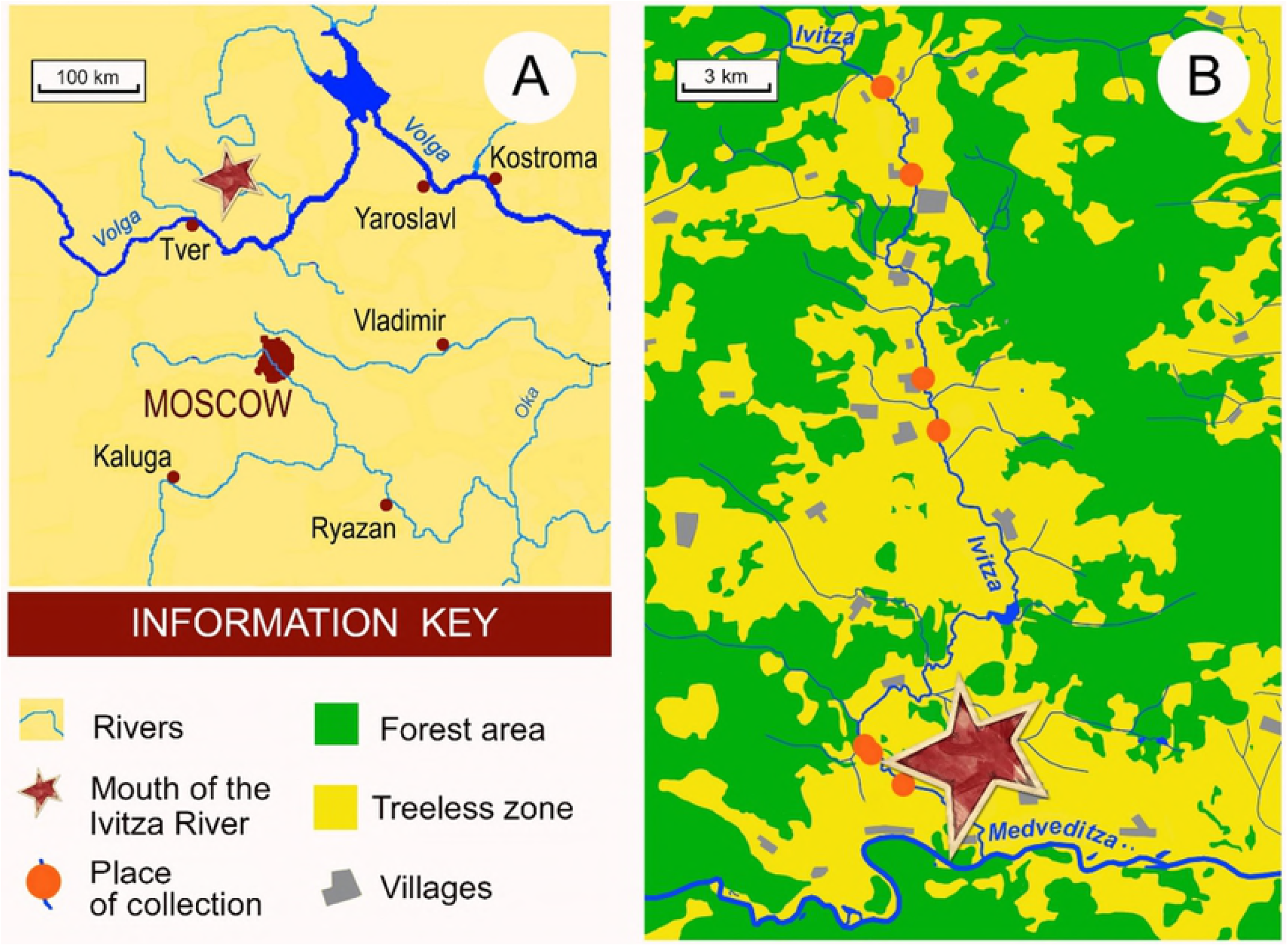
Collection area. A – Upper Volga River Basin. B – Places of specimen collection in the Ivitza River.

Specimens of freshwater Unionidae mussels were collected from 6 localities (Figure 1). From 1500 collected specimens 213 were selected for taxonomy analysis and the rest were returned to nature. Members of Unionidae are abundant in North-Eastern Europe and are not listed as endangered species therefore collection of specimens did not require any special permits. For the selected specimens a small piece of muscle foot tissue was placed in 95% ethanol (replaced next day with fresh one) and stored at −20°C.

All shells of the collected specimens are stored in the collection Federal Scientific Center of the East Asia Terrestrial Biodiversity, Far Eastern Branch of the Russian Academy of Sciences, Vladivostok, Russia.

### Identification of specimens by the modified comparative method

Identification of specimens was performed by V. Bogatov by means of the MCM. For the drawing of AMCVO, the stereoscopic binocular microscope (MBS-2) and Abbe’s drawing camera were used. The left valve was fixed with plasticine under the microscope objective with the front edge up in order to put polar optical axis of the microscope lay not only in its commissural plane, but also in the commissural planes of the earliest formation time (similar planes are outlined by the growth lines). For these purpose, the valve from its longitudinal axis (Fig 2.*1*) was deflected back until the field behind the keel disappeared from the field of view and fixed when the growth lines diverging from the umbo took the form of almost straight lines (Fig 2.*2*, 2.*3*). In this position, visible in the microscope contour of AMCVO in the projection onto the commissural plane/lateral surface of the shell should be almost straight or slightly curved toward the rear edge line with the angle to the longitudinal axis of the shell (Fig 2.*1*).

**Fig 2.**
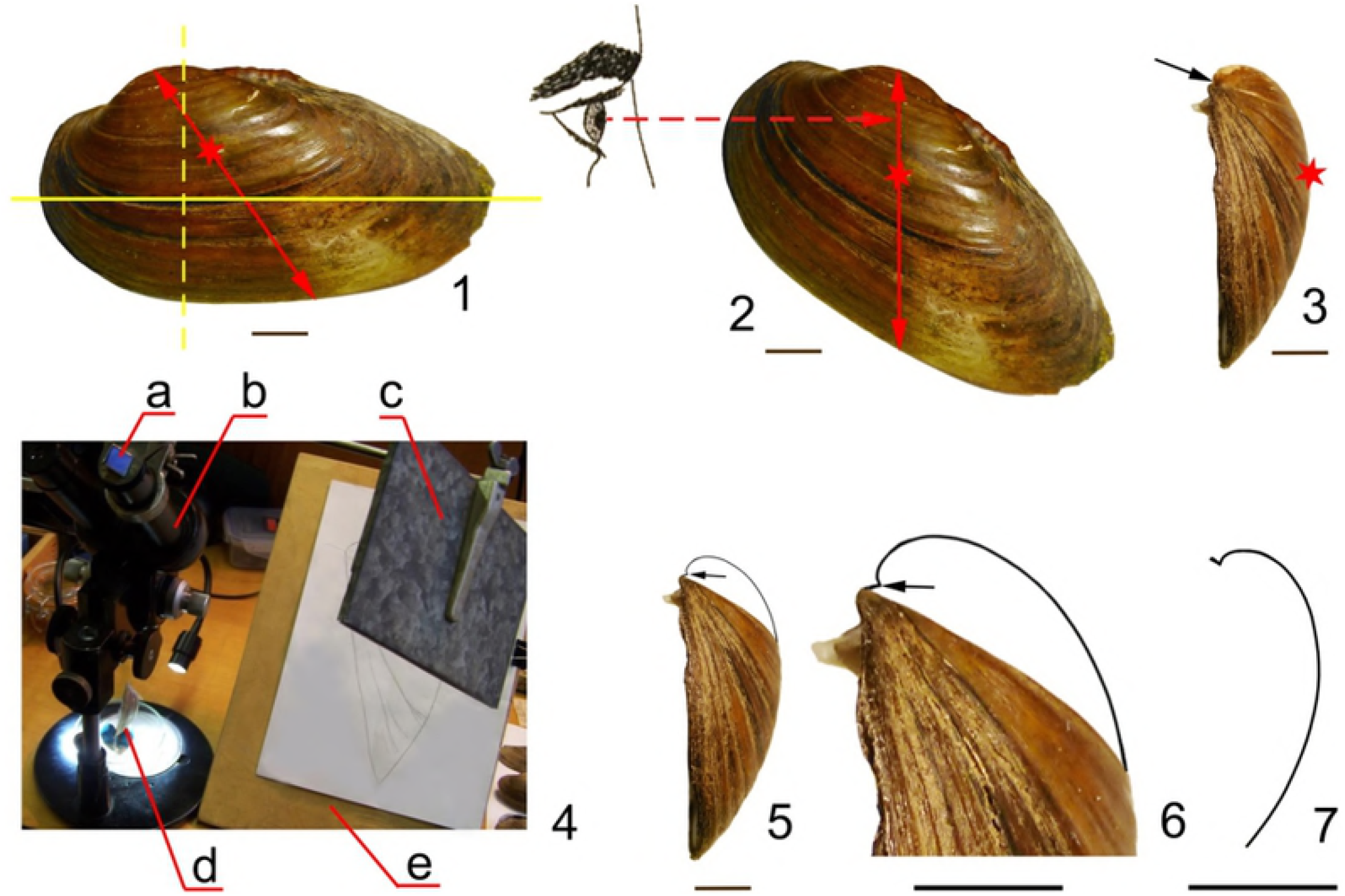
Identification of the contour of maximum convex of left valves of Unio (U-13) shells by the MCM. 1 - the side view of the shell; 2 - the position of the shell for the AMCVO drawing; 3 - the front view for the AMCVO drawing; 4 - stereoscopic binocular microscope mbs-2 with Abbe’s drawing camera (a - beam splitting prism, b - microscope, c - mirror, d - installed valve on a plasticine for viewing, e -sheet of paper for sketching outlines); 5 - plot of the upper third of AMCVO, 6 - the same at higher magnification; 7 - the final drawing of AMCVO. The yellow solid line through the valve shows the longitudinal axis of the shell, the yellow dashed line shows frontal section of the shell and the red line shows the lateral position of the AMCVO line, the red dashed line indicates the viewing direction for the AMCVO sketch, the asterisk indicates the most prominent point of the side surface of valve, the black arrows indicate the polar center of the spiral (the point from which the spiral shape turns). The scale bar is 1 cm.

The AMCVOs were observed at the magnification of 6×0.6 with drawing device (Fig 2.*4*), and the visible in the microscope upper part of contour was drawn (Figs 2.*5*, 2.*6*). The convergence point of the shell growth lines was taken as the initial point of deployment of the spiral contour (the polar center of the spiral) (Figs 2.*3*, 2.*5*, 2.*6*). To assess the reliability of the form by AMCVO, several drawings of the same object were performed, and the shell was installed under the microscope newly before each new drawing. Additionally, in the complex cases (severe corrosion of the shell, growth defects, etc), the contours of the left and the right valves of the same shell were compared, then the left valve was compared with the inverted picture of the right valve (Fig 2.*7*). If the contours of these figures matched, the resulting shape of the curve was considered as reliable. The contours of maximum convex of species identified by modified comparative method were compared with similar contours of shells from the systematic collection of mollusks stored in the Zoological Institute of the Russian Academy of Sciences (St. Petersburg). In this case drawing of AMCVOs was performed with the same drawing device (Fig 2.*4*).

### Genetic analysis

For 70 specimens of Unionidae mussels (Table 4) genomic DNA was extracted from foot tissues using the Diatom DNA Prep 200 reagents kit (“Laboratoriya Isogen” LLC, Russia) according to the manufacturer’s protocol. The designed primers (Table 2) were used to amplify the internal transcribed spacer 1 (ITS1) region, mitochondrial cytochrome oxidase gene (COI) and 16S rDNA region. All PCR reactions were performed in 25 μL volumes from 200 ng of the total DNA with 10 pmol of each primer and the PCR ScreenMix-HS (“Evrogene” Uppsala, Sweden). Thermocycling included one cycle at 95°C (5 min), followed by 34 cycles of 95°C (15 sec), 50-55°C (30 sec), and 72°C (45 sec) and a final extension at 72°C (5 min). Forward and reverse sequencing was performed on an automatic sequencer (ABI PRISM 3730, Applied Biosystems) using the ABI PRISM BigDye Terminator v. 3.1 reagent kit from Applied Biosystems (CA 94494, USA), and a cycling profile of 25 cycles of 96 °C for 30 s, 50 °C for 30 s, and 60 °C for 4 min. In addition, 19 sequences were obtained from NCBI’s GenBank, including sequences of *Margaritifera margaritifera* (L. 1758) as an outgroup for phylogenetic analysis (Table 3).

**Table 2.**
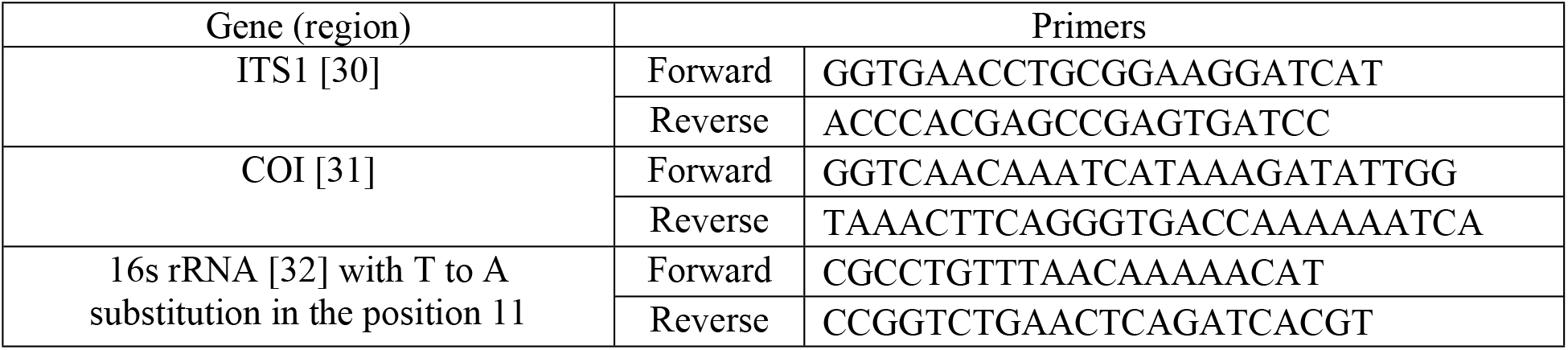
Primers for ITS1 region, the mitochondrial COI gene and 16s RNA region used for genetic analysis

**Table 3.** GenBank sequences included in analysis

| Species | Gene (region) | GenBank ID |
| --- | --- | --- |
| *Anodonta anatina* | ITS1 | DQ060178 |
|  | COI | KF030968 |
|  | 16S | KF030967 |
| *Anodonta cygnea* | ITS1 | DQ060185 |
|  | COI | KX822633 |
|  | 16S | DQ060164 |
| *Pseudanodonta complanata* | ITS1 | DQ060182 |
|  | COI | DQ060173 |
|  | 16S | DQ060166 |
| *Unio pictorum* | ITS1 | DQ060189<br>AJ295291 |
|  | COI | KC429109 |
|  | 16S | KC429266 |
| *Unio tumidus* | ITS1 | DQ060191 |
|  | COI | GU230750 |
|  | 16S | JQ253853 |
| *Unio crassus* | ITS1 | KJ525964 |
|  | COI | JX046553 |
|  | 16S | DQ060162 |
| *Unio cf. courtilieri* | COI | KC703900 |
| Genus *Margaritifera*<br>(outgroup for phylogenic<br>analysis) | ITS1 | DQ060193 |
|  | COI | DQ060171 |
|  | 16S | DQ060167 |

All sequences were aligned with ClustalW algorithm implemented in MEGA7 with default settings [33]. Phylogenic trees were built using maximum likelihood algorithm implemented in MEGA7 with 500 bootstrap replicates. Missed information or gaps were treated as complete deletion.

## Results

### Taxonomic analysis with modified comparative method

Analysis of the contours of maximum convex of 213 selected specimens with MCM identified five groups of comparative species: four genera (*Colletopterum, Pseudanodonta, Crassiana* and *Unio*) and two subgenera *Unio* (*Unio* s. str. and *Tumidiana*) according to Starobogatov-Bogatov system (Table 1) [15]. Fourteen comparative species of Unionidae mussels were identified (Figs 3, 4, and 5, Table 4). In the Ivitza River, we did not register *Anodonta* and *U. (U.) limosus* Nilsson 1822 specimens previously reported in the North-Eastern Europe [15]. The AMCVOs of *C. anatinum* and *C. ponderosum* showed no difference comparing to specimens from the collection of RAS Zoological Institute identified by Zhadin (Fig 3, *1a*). However *C. anatinum* was well distinguished from *C. ponderosum* by the umbo shifted strongly to the front edge of the shell and oval shell shape, while *C. ponderosum* shell shape is elongated [2] (Figs 4, *1a, 1b*).

**Fig 3.**
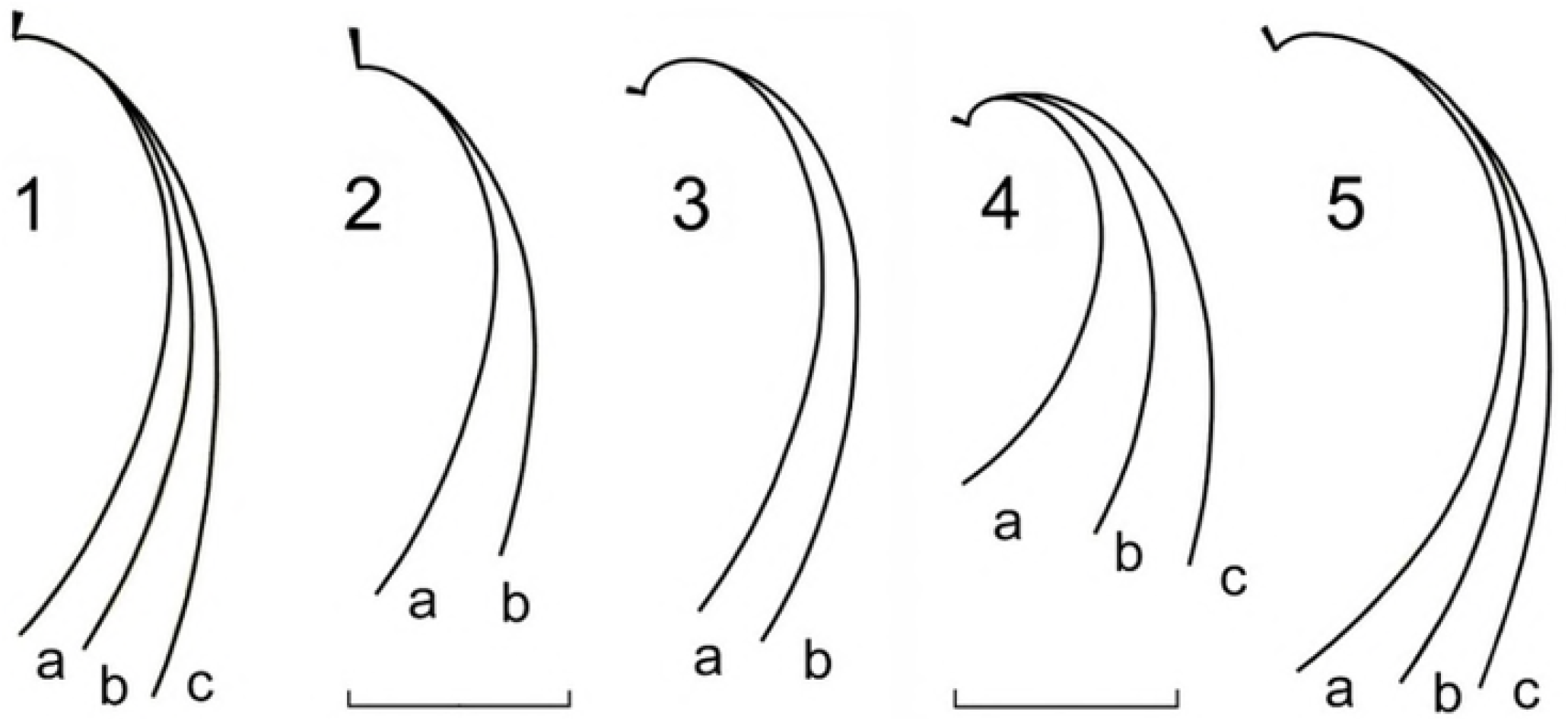
The contours of maximum convex of left valves of shells from the investigated species. Genus Colletopterum: 1a – *C. ponderosum* and *C. anatinum*, 1b – *C. piscinale*, 1c – *C. nilssoni*, genus *Pseudanodonta:* 2a – *P. elongata*, 2b – *P. complanata*, genus *Unio*: 3a – *U. pictorum*, 3b – *U. protractus*, 4a – *U. conus*, 4b – *U. longirostris*, 4c – *U. tumidus*, genus *Crassiana:* 5a – *Cr. crassa*, 5b – *Cr. musiva*, 5c – *Cr. nana*. Scale is 1 cm.

**Fig 4.**
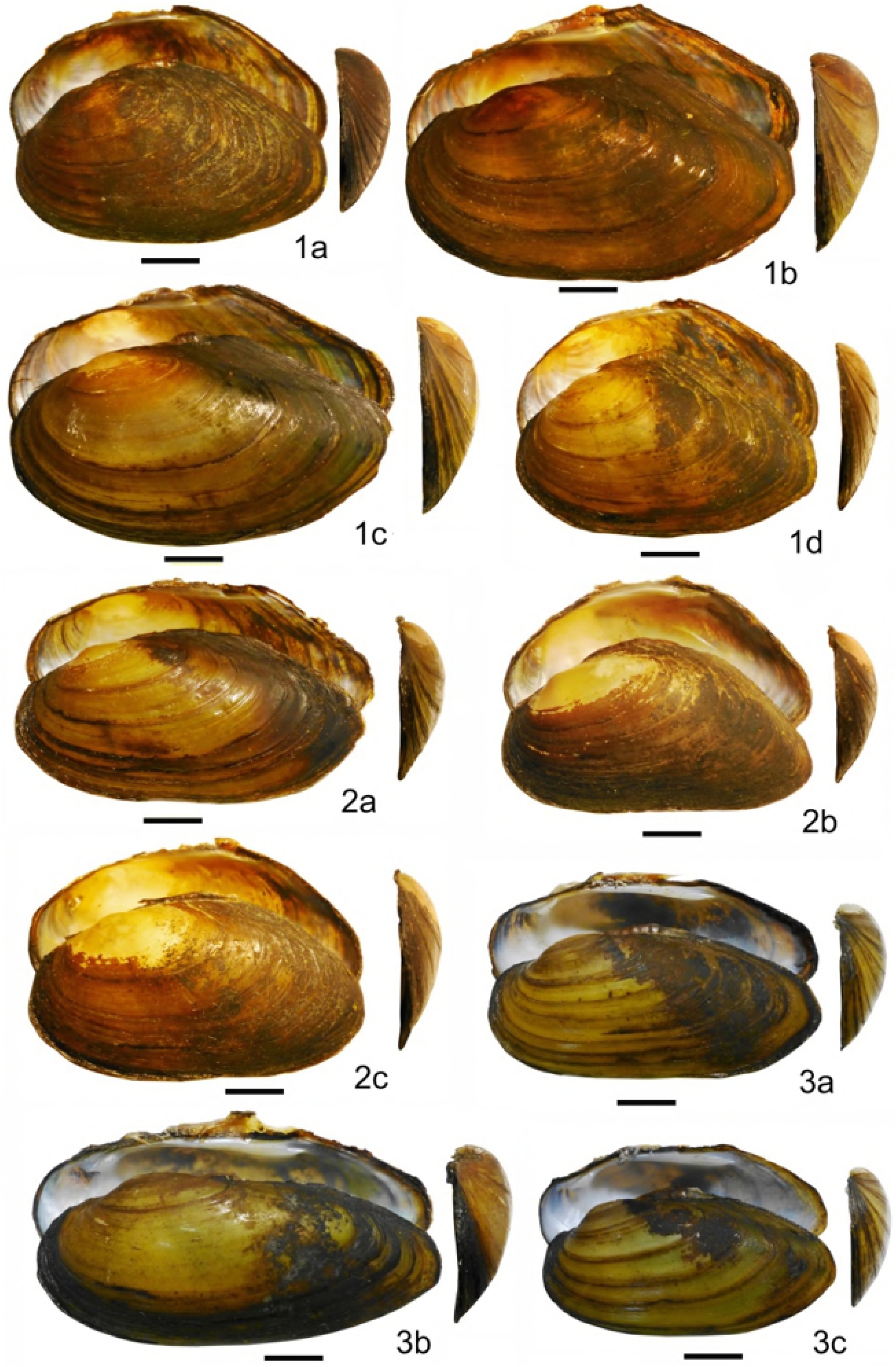
Shells of investigated Unionidae mussels of *Colletopterum, Pseudanodonta* and *Unio* genera (side, inside view and frontal view of the left valve). 1a – *C. anatinum*, 1b – *C. ponderosum*, 1c – *C. piscinale*, 1d – *C. nilssoni*. 2a – *P. elongata*, 2b – *P. elongata* (shortened form), 2c – *P. complanata*, 3a – *U. pictorum* (form with insufficient growth), 3b – *U. pictorum*, 3c – *U. protractus*. Scale is 1 cm.

**Fig 5.**
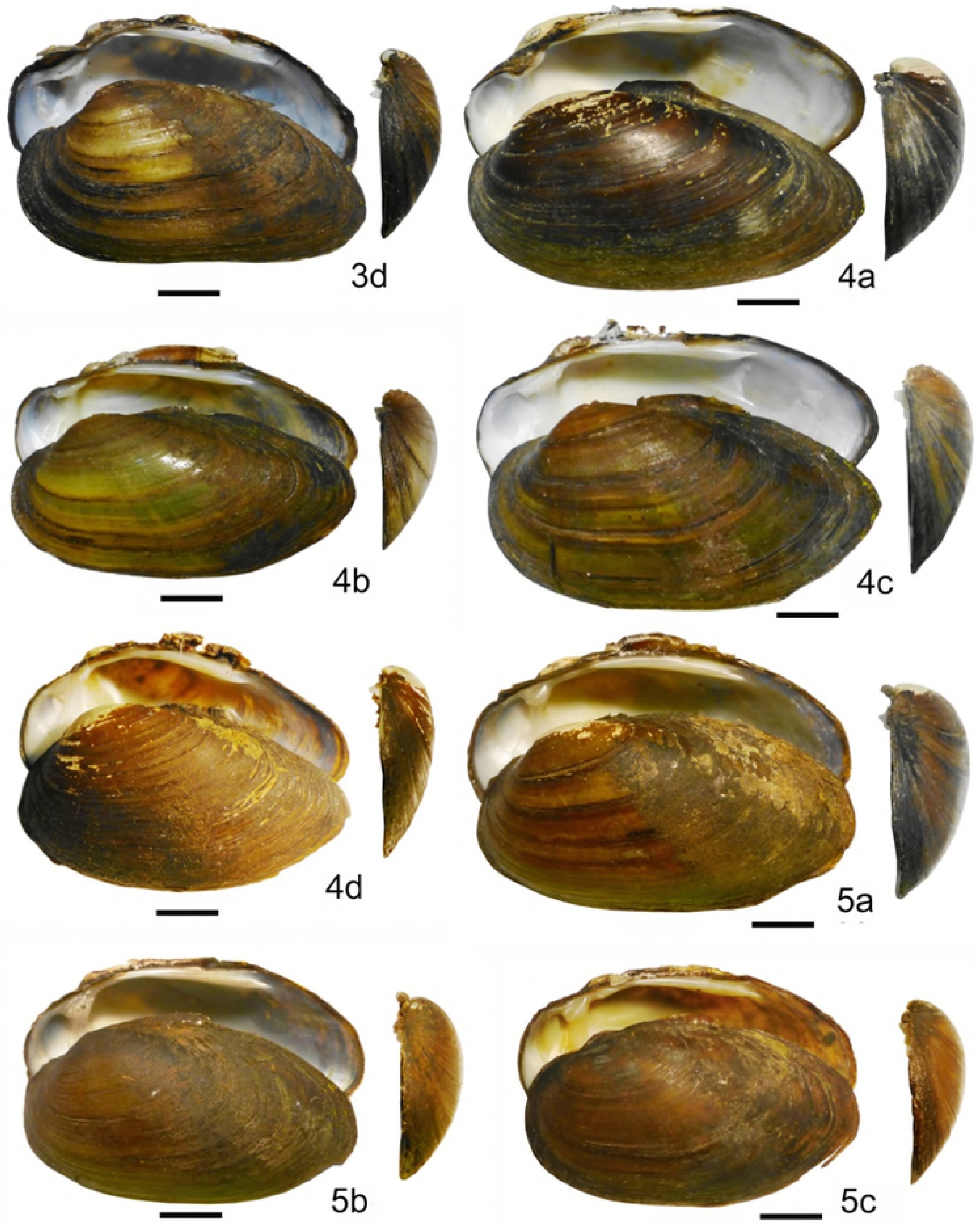
Shells of investigated Unionidae mussels of *Unio* and *Crassiana* genera (side, inside view and frontal view of the left valve). 3d – *U. protractus* (shortened form), 4a – *U. conus*, 4b – *U. longirostris* (elongated), 4c – *U. longirostris*, 4d – *U. tumidus*, 5a – *Cr. crassa*, 5b – *Cr. musiva*, 5c – *Cr. nana*. Scale is 1 cm.

**Table 4.**
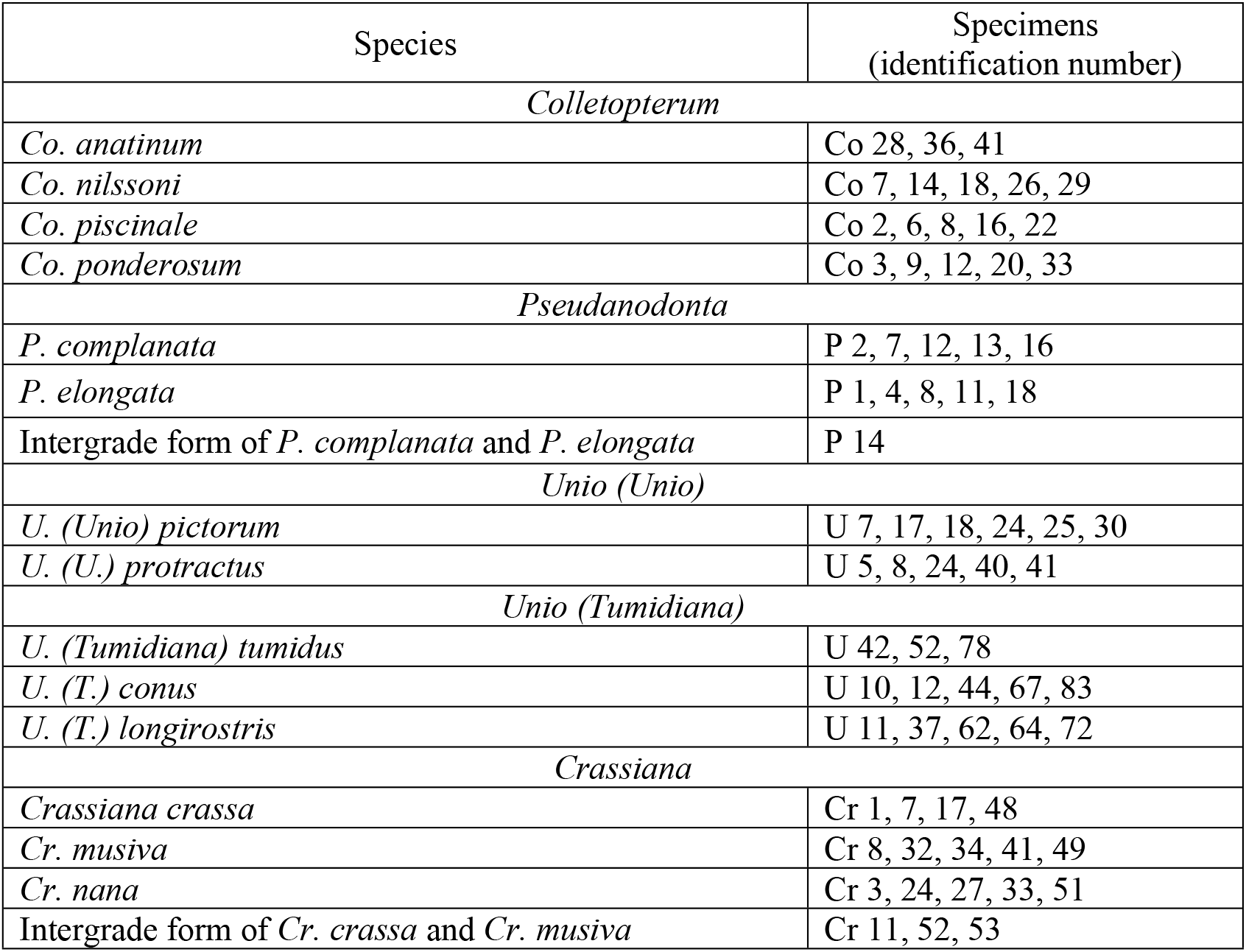
Taxonomic composition of Unionidae from Ivitza River based on Starobogatov-Bogatov system [15] and list of specimens selected for genetic analysis

| Species | Specimens<br>(identification number) |
| --- | --- |
| *Colletopterum* |  |
| *Co. anatinum* | Co 28, 36, 41 |
| *Co. nilssoni* | Co 7, 14, 18, 26, 29 |
| *Co. piscinale* | Co 2, 6, 8, 16, 22 |
| *Co. ponderosum* | Co 3, 9, 12, 20, 33 |
| *Pseudanodonta* |  |
| *P. complanata* | P 2, 7, 12, 13, 16 |
| *P. elongata* | P 1, 4, 8, 11, 18 |
| Intergrade form of *P. complanata* and *P. elongata* | P 14 |
| *Unio (Unio)* |  |
| *U. (Unio) pictorum* | U 7, 17, 18, 24, 25, 30 |
| *U. (U.) protractus* | U 5, 8, 24, 40, 41 |
| *Unio (Tumidiana)* |  |
| *U. (Tumidiana) tumidus* | U 42, 52, 78 |
| *U. (T.) conus* | U 10, 12, 44, 67, 83 |
| *U. (T.) longirostris* | U 11, 37, 62, 64, 72 |
| *Crassiana* |  |
| *Crassiana crassa* | Cr 1, 7, 17, 48 |
| *Cr. musiva* | Cr 8, 32, 34, 41, 49 |
| *Cr. nana* | Cr 3, 24, 27, 33, 51 |
| Intergrade form of *Cr. crassa* and *Cr. musiva* | Cr 11, 52, 53 |

The AMCVOs of one *Pseudanodonta* and three *Crassiana* shells were different from the contours defined for the species and were conditionally classified as intergrade forms (Table 4). It should be noted that in some groups of comparative species were found shells with insufficient growth (e.g. Fig 4, *3a*), elongated form (Fig 5, *4b*) or shortened rear part, because of the lag in the growth of the valves rear part (e.g. Fig 4, *2b*; Fig 5, *3d*).

### Taxonomic analysis with molecular genetic markers

From 213 collected specimens of Unionidae mussels, 70 were analyzed by genetic methods. We investigated species variation by comparing internal transcribed spacer 1 (ITS1) sequence in the nuclear ribosomal repeat DNA region with those of two of the most commonly used genes for the phylogenetic analysis in the group, mitochondrial cytochrome oxidase (COI) and 16S ribosomal DNA (16S rDNA). The ITS1 region, 16S rDNA and COI gene were sequenced from 60, 68 and 19 samples, respectively (Table 5). GenBank IDs of all sequences obtained in this study are indicated in Table S1. Mitochondrial gene regions are common genetic markers in Unionidae taxonomic analysis [18,21]. We complemented our study with ITS1 region evaluated in previous studies as an appropriate genetic marker for combination with mitochondrial genes in phylogenetic analysis of Unionidae [26,27].

**Table 5.** Lengths of the ITS1, 16S rDNA and COI regions obtained for the investigated genera.

| Gene (region) | Genus | Samples | Lengths (bp) of region |
| --- | --- | --- | --- |
| ITS1 | *Colletopterum* | Co 3, 6, 7, 8, 9, 12, 16, 18, 22, 26, 29, 33 | 539 |
|  | *Pseudanodonta* | P 1, 2, 4, 7, 8, 11, 12, 13, 14, 16, 18 | 536–561 |
|  | *Unio (Unio)* | U 5, 7, 8, 17, 18, 24, 25, 30, 40, 41 | 299 |
|  | *Unio (Tumidiana)* | U 10, 11, 12, 37, 44, 62, 64, 67, 72, 78, 83 | 381–520 |
|  | *Crassiana* | Cr 1, 3, 7, 8, 11, 17, 24, 27, 32, 33, 34, 41, 49, 51, 52, 53 | 377–489 |
| 16S rDNA | *Colletopterum* | Co 2, 3, 6, 7, 8, 9, 12, 14, 16, 18, 20, 22, 26, 28, 29, 36, 41 | 530–545 |
|  | *Pseudanodonta* | P 1, 2, 4, 7, 8, 11, 12, 13, 14, 16, 18 | 363 |
|  | *Unio (Unio)* | U 5, 7, 8, 17, 18, 24, 25, 30, 40, 41 | 385–542 |
|  | *Unio (Tumidiana)* | U 10, 11, 12, 37, 42, 44, 52, 62, 64, 67, 72, 78, 83 | 390–543 |
|  | *Crassiana* | Cr 1, 3, 7, 8, 11, 17, 24, 27, 32, 33, 34, 41, 48, 49, 51, 52, 53 | 372–546 |
| COI | *Colletopterum* | Co 2, 6, 12, 20, 41 | 295 |
|  | *Pseudanodonta* | P 8, 13, 16, 18 | 295 |
|  | *Unio (Unio)* | U 5, 18, 30, 41 | 295 |
|  | *Unio (Tumidiana)* | U 42, 44 | 295 |
|  | *Crassiana* | Cr 8, 32, 33, 48 | 295 |

Maximum likelihood phylogenic tree was constructed for 66 sequences (60 samples and six GenBank sequences including *M. margaritifera* as an outgroup). Combined multiple alignments of ITS1 and 16S sequences was 602 bases long with 257 bp for ITS1 and 345 bp for 16S rDNA. Investigated samples were divided into the five groups with 98-100% bootstrap support (Fig 6). Each group contained different species identified by morphological criteria and represented comparative genera *Pseudanodonta, Crassiana, Colletopterum* and two subgenera of genus *Unio* – *Unio* s. str. and *Tumidiana*. Sequences presented in one group with 98-100% support are likely to belong to one species calling morphological criteria into question.

**Fig 6.**
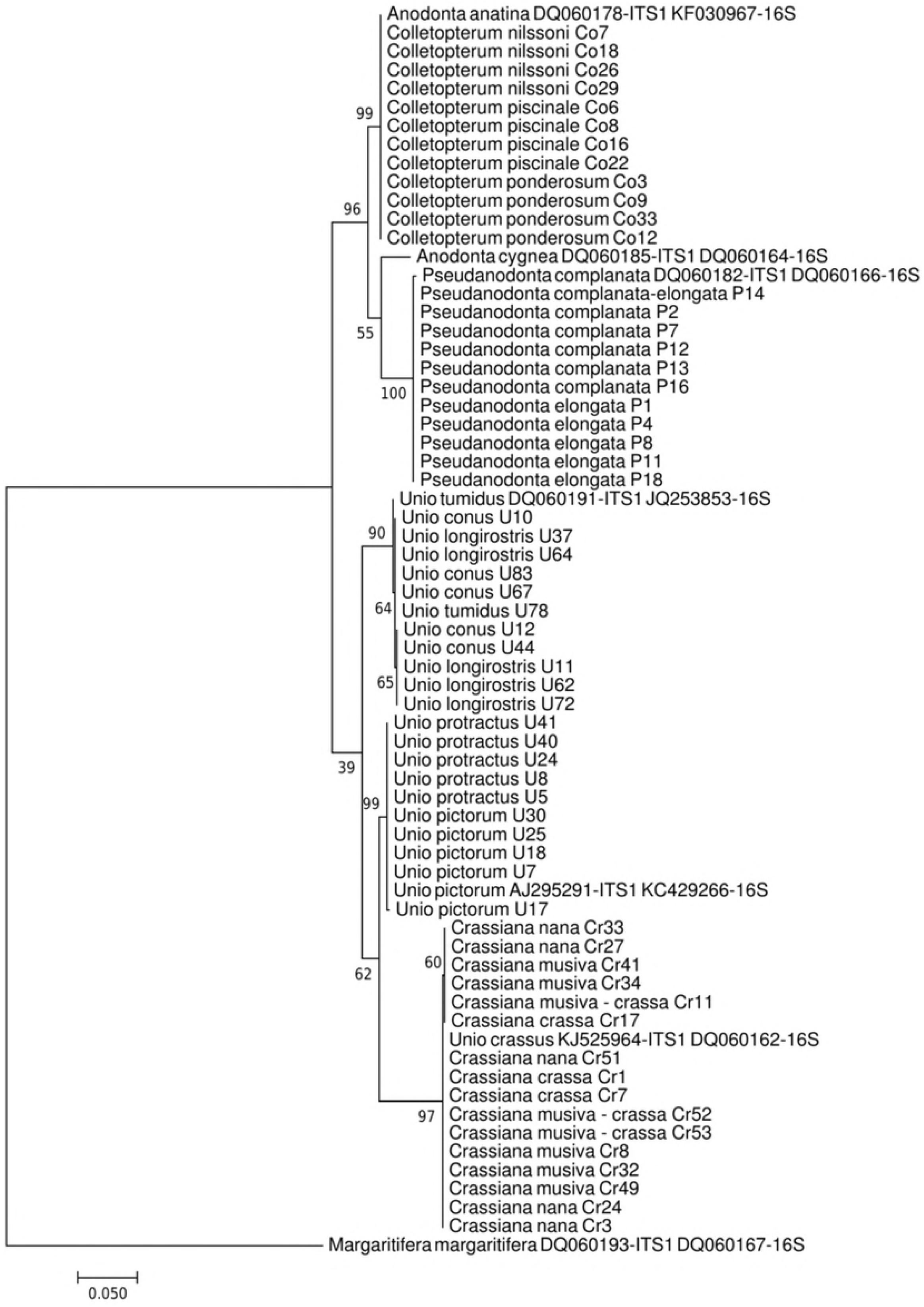
Maximum likelihood phylogenic tree for 602 bases long combined multiple alignment of ITS1 (257 bp) and 16S (345 bp) sequences with bootstrap 500 support values indicated at nodes. Missed information or gaps were treated as complete deletion. *M. margaritifera* was used as an outgroup.

Another phylogenetic tree was built for COI 295 bases long alignment (Fig 7) because the smaller set of sequences was obtained for COI than for ITS1 and 16S regions. The same five groups were observed as for ITS1 and 16S combination tree although some differences in relationships between groups were shown.

**Fig 7.**
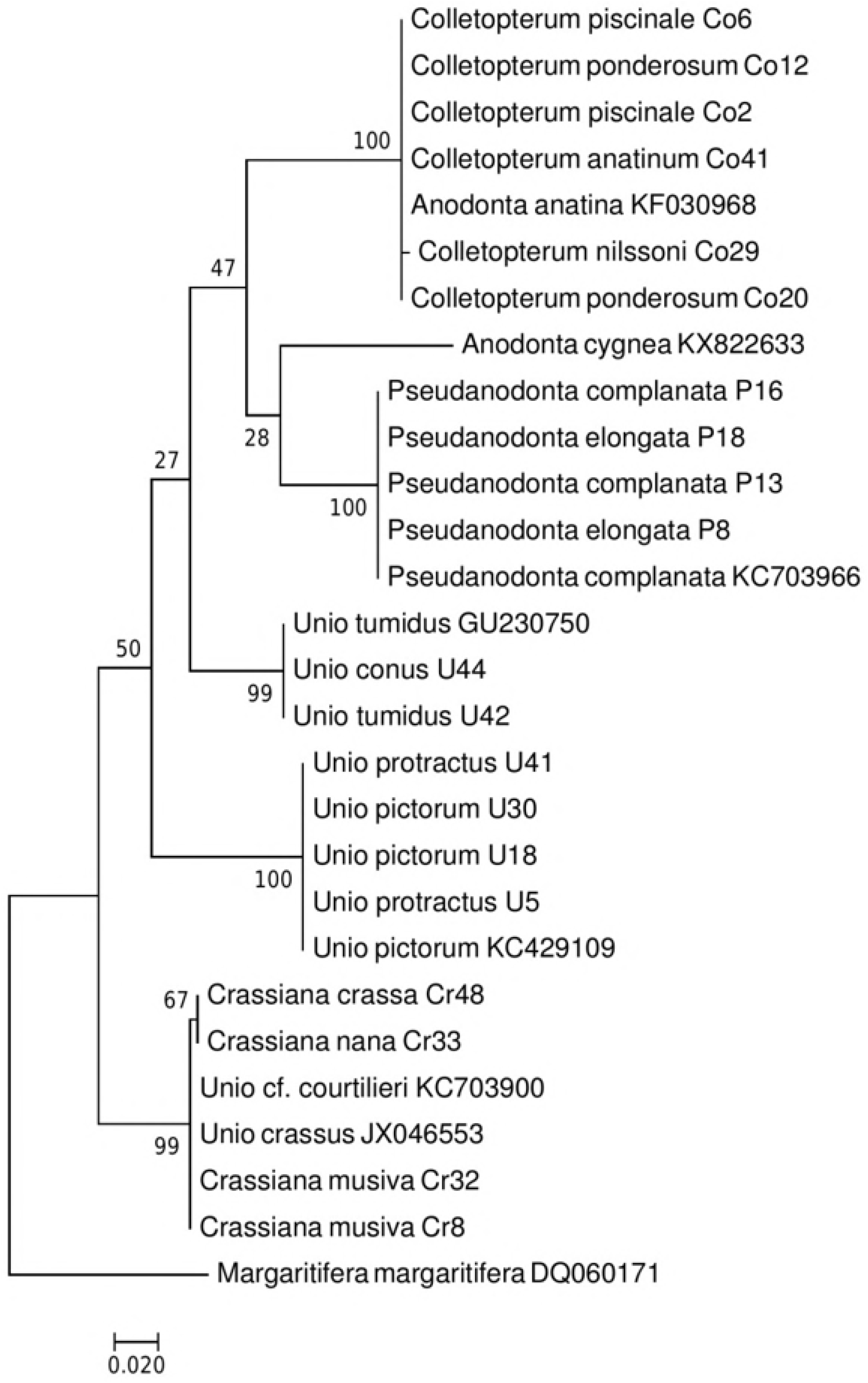
Maximum likelihood phylogenic tree for 295 bases long multiple alignment of COI sequences with bootstrap 500 support values indicated at nodes. Missed information or gaps were treated as complete deletion. *M. margaritifera* was used as an outgroup.

For estimation of variation within and between groups, we calculated P distances for ITS1, 16S rDNA and COI multiple alignments including GenBank sequences presented in table 3 by MEGA7. P distance is the number of base differences per site from averaging over all sequence pairs between and within groups with all ambiguous positions removed for each sequence pair. Within-group and between-group variation is shown in Table 6. Within-group variation was small (< 1%) comparing to between-group variation indicating small differences between species identified by morphological criteria.

**Table 6.** Estimated p distances for ITS1, COI and 16S rDNA gene regions.

| ITS1 region (68 nucleotide sequences with 528 positions) |  |  |  |  |  |
| --- | --- | --- | --- | --- | --- |
|  | *Colletopterum* | *Pseudanodonta* | *Unio (Unio)* | *Unio (Tumidiana)* | *Crassiana* |
| *Colletopterum* | 0 |  |  |  |  |
| *Pseudanodonta* | 0.20032 | 0 |  |  |  |
| *Unio (Unio)* | 0.05649 | 0.09728 | 0.00035 |  |  |
| *Unio (Tumidiana)* | 0.19728 | 0.11844 | 0.05044 | 0.00284 |  |
| *Crassiana* | 0.1789 | 0.11183 | 0.06523 | 0.08185 | 0.00067 |
| *M. margaritifera* | 0.60289 | 0.60051 | 0.59766 | 0.60661 | 0.59487 |
| COI gene region (27 nucleotide sequences with 259 positions) |  |  |  |  |  |
|  | *Colletopterum* | *Pseudanodonta* | *Unio (Unio)* | *Unio (Tumidiana)* | *Crassiana* |
| *Colletopterum* | 0.00097 |  |  |  |  |
| *Pseudanodonta* | 0.10557 | 0 |  |  |  |
| *Unio (Unio)* | 0.14076 | 0.11695 | 0.03842 |  |  |
| *Unio (Tumidiana)* | 0.12155 | 0.09831 | 0.09944 | 0 |  |
| *Crassiana* | 0.13646 | 0.10983 | 0.09514 | 0.10644 | 0.00203 |
| *M. margaritifera* | 0.18305 | 0.15593 | 0.16497 | 0.14915 | 0.14373 |
| 16S rDNA region (75 nucleotide sequences with 523 positions) |  |  |  |  |  |
|  | *Colletopterum* | *Pseudanodonta* | *Unio (Unio)* | *Unio (Tumidiana)* | *Crassiana* |
| *Colletopterum* | 0.00042 |  |  |  |  |
| *Pseudanodonta* | 0.06061 | 0 |  |  |  |
| *Unio (Unio)* | 0.09538 | 0.12343 | 0.00052 |  |  |
| *Unio (Tumidiana)* | 0.10175 | 0.10497 | 0.07294 | 0.00074 |  |
| *Crassiana* | 0.11897 | 0.13116 | 0.04759 | 0.08089 | 0.00135 |
| *M. margaritifera* | 0.21718 | 0.27348 | 0.25648 | 0.22935 | 0.24379 |

In the result of molecular genetic analysis, genetic homogeneity in all five groups of the comparative species of the North-Eastern European Unionidae was identified. Taken together, morphological and genetic analysis of taxonomic composition of Unionidae from the Ivitza River revealed five species belonged to three genera *Anodonta*, *Pseudanodonta* and *Unio* (Table 7).

**Table 7.** Taxonomic composition of Unionidae mussels from the Ivitza River. Comparative species shown to be genetically homogeneous are indicated in parentheses

| Class Bivalvia |  |
| --- | --- |
| Family Unionidae |  |
| Genus | Species |
| *Anodonta* (= *Colletopterum*) | *A. anatina* (= *C. ponderosum*, *C. piscinale*, *C. nilssoni*) |
| *Pseudanodonta* | *P. complanata* (= *P. elongata*) |
| *Unio* (= *Crassiana*) | *U. pictorum* (= *U. protractus*) |
|  | *U. tumidus* (= *U. longirostris*, *U. conus*) |
|  | *U. crassus* (= *Cr. musiva*, *Cr. nana*) |

## Discussion

We employed modified comparative method (MCM) and methods of molecular genetics to analyze a large, comprehensive sample of Unionidae mussels from the Ivitza River. MCM attributes 213 individuals from this sample to 14 morphological, or comparative, species belonging to five groups according to their AMCVOs (Table 1, Fig 3). AMCVOs of only four samples are different from AMCVOs established for comparative species presumably because of umbo deformation (Table 4).

However, analysis of nuclear and mitochondrial molecular markers subdivides the 213 individuals into only 5 highly homogeneous groups, each of which very likely represents a separate population. Substantial genetic distances between these groups (Figs 6, 7) leave little doubt that each of them represents a separate biological species. Data on mitochondrial and nuclear genes are in perfect agreement with each other.

Therefore, we can conclude that comparative method often attributes to different species individuals that belong to the same population of one species. This must be a result of a wide morphological variation within species of Unionidae and is consistent with previous works on *Unio* [18] and *Colletopterum* taxonomy [21].

As far as the group *Pseudanodonta* (priority name *Ps. complanata*) is concerned, which is more phylogenetically distant from *A. cygnea*, we cannot recommend to treat it as a synonym of *Anadonta*. A number of authors [34–36] also treat *Pseudanadonta* as a separate genus. However, because *Pseudanodonta* is rather close genetically to *A. cygnea* and *A. anatina*, there is no reason to attribute this genus to the subfamily Pseudanodontinae, as it has been done before [7,15]. Instead, the genus *Pseudanodonta* should be transferred into the subfamily Anodontinae according to Bogatov and Kijashko [15], or into the subfamily Unioninae (tribe Anodontini) according to Carter et al. [37], and the name Pseudanodontinae should be treated as invalid.

At the same time, we cannot currently agree with Klishko et al. [21] who proposed to merge the nominative subgenus *Colletopterum* (type species: *Anodonta letourneuxi* Bourguignat 1880 = *Anodonta-subcircularis* Clessin 1873) with *A. anatina*. Indeed, so far not a single definite individual from the subgenus *Colletopterum* has been studied genetically. Morphologically, mollusks that were attributed to the comparative subgenera *Piscinaliana* Bourguignat 1881 (type species: *Colletopterum piscinale*) and *Colletopterum* are substantially different from each other [13,15], so that it is entirely possible that the comparative subgenus *Colletopterum* represents a distinct species. So far, this group of bivalves remains the least studied one among the European Unionidae.

Short phylogenetic distances between individuals attributed to *Colletopterum* by MCM and the type species of genus *Anodonta, A. cygnea* (Figs 6, 7), obtained from GenBank, implies that all these individuals, which were previously included into subgenus *Piscinaliana* [15], must instead be included into genus *Anodonta*. According to the principle of priority, all the forms from this group that were recognized by MCM should be called *Anodonta anatina*. As a result, there is also no need to subdivide the genus *Colletopterum* into subgenera *Colletopterum* s. str. and *Piscinaliana*. Taxonomic position of *A. anatina* within genus *Anodonta* has been acknowledged before [38,39].

For genetic groups that belong to the subfamily Unioninae (tribe Unionini), the following names have priority: *U. pictorum* for the group of для *U. pictorum, U. limosus* and *U. protractus*, and *U. tumidus* for the group of *U. tumidus, U. longirostris* and *U. conus. U. tumidus* definitely should be included into genus *Unio*, despite it being attributed to the genus *Tumidiana* (type species: *Unio tumidus)*, in early classification of the comparative species [7] (Table 1). In contrast, caution is advised regarding transferring members of *Crassiana* (priority name *U. crassus*) into the genus *Unio*, as suggested by [35,36]. Still, there is no reason to retain *U. crassus* within the subfamily Psilunioninae (type species: *Psilunio* Stefanescu 1896) (Table 1). We agree with Klishko et al. [18] that, in order to determine the taxonomical position of *Unio crassus* further molecular studies are necessary.

Therefore, Unionidae in the river Ivitza are represented by five species which belong to two tribes and three genera in Unioninae subfamily: tribe Anodontini, genus *Anodonta* – *A. anatina* (= *Colletopterum ponderosum, C. piscinale, C. nilssoni);* genus *Pseudanodonta* – *Ps. complanata* (= *Ps. elongata);* tribe Unionini, genus *Unio* – *U. pictorum* (= *U. protractus), U. tumidus* (= *U. longirostris, U. conus*), and *U. crassus* (= *Crassiana musiva, Cr. nana*).

Our data indicate that freshwater Unionidae possess high phenotypic plasticity, manifested in their intraspecific variation of morphological characters. On top of AMCVO, other traits that were previously used to define the so-called biological species [2] are also highly plastic. In particular, the general shape of the shell, which was considered to be a species-defining trait, can vary in Unionini from ovoid or sphenoid to elongated oval (as in U. *tumidus)*. In Anodontini, the same trait can vary from oval or rounded triangular to elongated oval (as in *A. anatina)*. Similarly, the shape of the front edge of the shell (such as narrow, as in *A*. *anatina* or wide, as in *A. ponderosum*., according to the Zhadin’s key) lacks taxonomic significance. The same is true regarding the location of the tip of the shell relative to front edge of the valve, its thickness, etc. Also, in Bivalvia the growth of the rear edge of the shell may get faster or slower with age, which makes identification by the shell problematic. For example, the general shape of the shell of *U. pictorum* may become similar to the elongated shape of the shell of U. *tumidus* (Fig 5, *4b*) and, *vice versa*, truncated form of *U. pictorum* (Fig 5, *3d*) can be easily confused with the standard form of U. *tumidus* (Fig 5, *4d*). As a result, a substantial proportion of species and subspecies recognized by Zhadin [2] is not supported by the modern taxonomical systems.

Within established comparative groups of Bivalvia mollusks, considered previously as comparative genera or species, there is only a rather limited number of forms (no more than three, and only rarely four [40,41]) with specific AMCVO, from the most convex to the most flat. This pattern remains obscure and needs to be studied further. In this case, as well as in addressing other issues related to the shell morphology, the comparative method is definitely useful, as it currently provides the best approach to characterizing the degree of convexity of their valves even in similar forms, being more efficient than using the ratios of the principal measurements of the shell.

Of course, a complete revision of Eastern European Unionidae, based on molecular data, would require sampling individuals from larger number of different locations. Still, our results strongly suggest that the number of their real species is much smaller than commonly assumed. Thus, comparative method (including the modified Bogatov’s method MCM) could not be applied for the species identification not only in investigating, but, apparently, in most other freshwater Unionidae.

## Acknowledgements

We thank I. E. Arakcheev for helping with samples collection on the Ivitza River and G. D. Kolbasova for tissue preparation. The work is being supported by Far Eastern Branch of Russian Academy of Sciences (grant number № 18-4-039) and by Russian Science Foundation grant N 16-14-10173.

## Supporting information

**Table S1. GenBank IDs of all obtained sequences of ITS1, 16S, COI regions of Unionidae mussels.**

